# Synchronous assembly of peptide anisosome

**DOI:** 10.1101/2024.07.07.602414

**Authors:** Laicheng Zhou, Longcheng Zhu, Cong Wang, Tengyan Xu, Jing Wang, Bin Zhang, Xin Zhang, Huaimin Wang

## Abstract

Biomolecular condensates, formed by liquid-liquid phase separation (LLPS) of proteins or the complex of protein and nucleic acids, play key roles in regulating physiological events in biological system. However, the formation of mono-component yet inhomogeneous condensates is limited, and the underlying mechanism remains unclear. Here, we report the symmetrical core-shell structural biomolecular condensates formed through the LLPS by programming a tetra-peptide library. Mechanistic studies reveal that the tryptophan (W) is critical for the formation of core-shell structure because of its stronger homotypical π-π interaction compared with other amino acids, which endow us to modulate the droplets from core-shell to homogeneous structures by encoding the amino acid composition. Using molecular dynamics (MD) simulation and molecular engineering, we find that the inner core of LLPS is composed of dynamic and reversible fibers surrounded by liquid-like shells, resulting in a stable core-shell LLPS. Furthermore, we could control the multiphasic droplet formation by an intrinsic redox reaction or post-translational modification of peptide through phosphorylation, which facilitates the rational design of synthetic LLPS with various applications on demand.

## Introduction

Cells evolve to form numerous membrane-enclosed organelles and membrane-less compartments to ensure complex cellular activities occur in a crowding environment. More evidence has shown that liquid-liquid phase separation (LLPS) plays a crucial role in forming membrane-less compartments consisting of proteins and nucleic acids^1–3^. Recent advances have unraveled one particular type of intracellular condensate that is not a homogeneous fluid but contains a sub-structure represented by stress granules, anisosome, and nucleolus, among many others^4–8^. The dynamic formation and dissociation of such sub-compartments inside the condensates facilitate the regulation of the complex cellular activities more precisely and spatiotemporally^3, 9, 10^.

Intrinsically disordered proteins (IDPs) or intrinsically disordered regions (IDRs) have been identified and used for constructing biomolecular condensates through LLPS *in vitro* to delineate the molecular mechanism of phase transition behaviors, physicochemical properties, and explore their potential applications^11–13^. The dynamic and multivalent weak noncovalent interactions, such as electrostatic interactions, hydrophobic interactions, hydrogen bonds, π-π stacking, and cation-π interactions, among the intermolecular or intramolecular, often work cooperatively to drive the formation of condensates through the LLPS^14–17^. Based on the deciphered knowledge about LLPS, several synthetic LLPS made from IDPs/ IDRs derivatives, polypeptides/RNA complex, and short peptide derivatives have been developed^18–25^. However, only limited reports of multiphasic synthetic condensates have been achieved using multi-components of proteins, peptides/RNAs complex, and polymers^26–29^. To the best of our knowledge, multiphasic droplets formed by a single component of short peptide have not been reported.

In this work, we describe the formation of peptide anisosome through the LLPS by programming a tetra-peptide library (Fig. 1). Under the external stimuli (pH or temperature), the single component of the peptide could synchronously self-assemble into liquid droplets with a core-shell structure. The outer shell of the droplets has better fluidity than the inner core. Mechanistic studies by IR, Raman, and molecular engineering suggested that the hydrophobic interaction, π-π stacking, and hydrogen bonds contribute to the droplet formation through LLPS in the tetra-peptide system. The amino acids mutation and atomic modification results indicate that tryptophan (W) is crucial for forming the core-shell structure. Molecular engineering combined with the MD simulation experiments showed that the stronger homotypical π-π interaction is the main driving force for the synchronous assembly of peptide anisosome (Fig. 1a), which maintained the fibrous structures in the core. Using chemical engineering to control the balance of interactions between peptide-peptide and peptide-water, we can control the droplet with either a homogeneous or core-shell structure (Fig. 1b). The transformation of the multiphasic droplet and homogeneous structure could be easily controlled by an intrinsic redox reaction. Meanwhile, the distribution of the guest molecules can also be controlled spatiotemporally, reminiscing the function of the nucleosome (Fig. 1d). This work provides a general modular platform for constructing multiphasic droplets with a single component by tuning the strength of π-π interaction among the driving force for LLPS.

**Fig 1.**
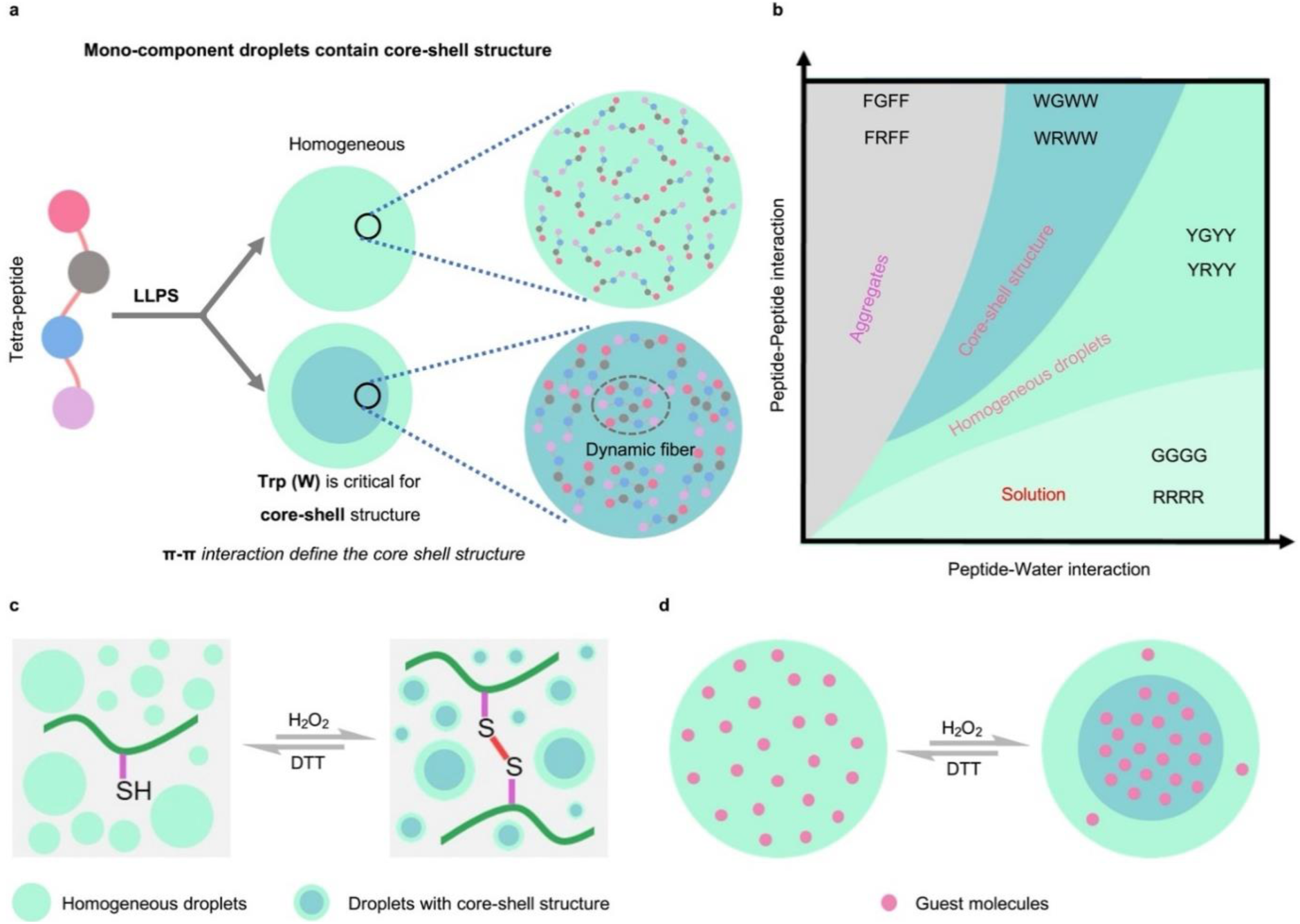
Liquid-liquid droplets with a core-shell structure formed by mono-component tetra-peptides. **a**, Schematic illustration of the microstructure of droplets that can be tuned by amino acid composition. **b**, Schematic illustration of the balance between peptide-peptide and peptide water interactions that define the phase transition behavior of tetra-peptides. **c**, Schematic illustration of the redox-responsive droplets. **d**, The homogeneously distributed guest molecules can be recruited into the newly formed inner core of droplets under the external stimuli.

## Results and discussion

### Design and the formation of droplets through LLPS

The molecular rules for phase separation of IDPs often involve polar residues. Tryptophan (W) rarely appears in IDPs and their contribution to phase separation is under appreciated. Associative polymers like proteins or polypeptides have higher valence than short peptides, enabling sufficient sticker-sticker interactions for phase separation^30–32^. By contrast, short peptides with limited valence require more enthalpy or entropy compensation to drive phase separation at a low concentration. Increasing the strength of site-specific interactions among short peptides could promote phase separation. Among the 20 amino acids, W has the largest hydrophobic and aromatic ring, providing the strongest π-system involved interactions and hydrophobic interactions, the NH group of the indole ring can further provide hydrogen bonds^33, 34^. Such strong interactions provided by W may cause the collapse of IDPs or IDRs. This factor partially contributes to why W presents a very limited ratio in the composition of both low complexity regions (LCRs) and intrinsically disordered regions (IDRs)^35^. In contrast, involving W seems to be the best choice to construct droplets of short peptides. Moreover, W exhibits distinctive and sharp Roman peaks, some of which has been demonstrated as a structural marker for the conformation, hydrogen bonding, hydrophobic interaction, and cation-π interaction, which could also facilitate our understanding of the molecular mechanisms for the formation of the liquid-liquid phase separation^36–38^.

We first designed and synthesized the peptide WRWW by taking advantage of the classical cation-π interactions between W and R^22, 33^. The N-terminus of the peptide is the amide group that can reduce their ability to self-assemble into hydrogels^23, 39^. Meanwhile, the peptide WGWW, which could not form cation-π interactions, was synthesized as a control group. Interestingly, both peptides can form liquid-like droplets, implying that the cation-π interactions between R and W may be unnecessary in our tetra-peptide system.

Next, we set a template-WXWW (X is the guest residue) and mutated the X with other 18 natural amino acids (Fig. 2a). Apart from 4 peptides (WVWW, WIWW, WDWW, and WEWW), the results showed that 16 peptides underwent liquid-liquid phase separation with different stability (Fig. 2c). Some of the droplets could transform into aggregates within a few minutes or hours. To comprehensively understand the properties and mechanism of the droplets, we focused on the 6 stable droplets (X = R, G, P, H, W, Y) that share a similar pH/temperature responsiveness (Fig. 2b, Fig. 2d).

**Fig 2.**
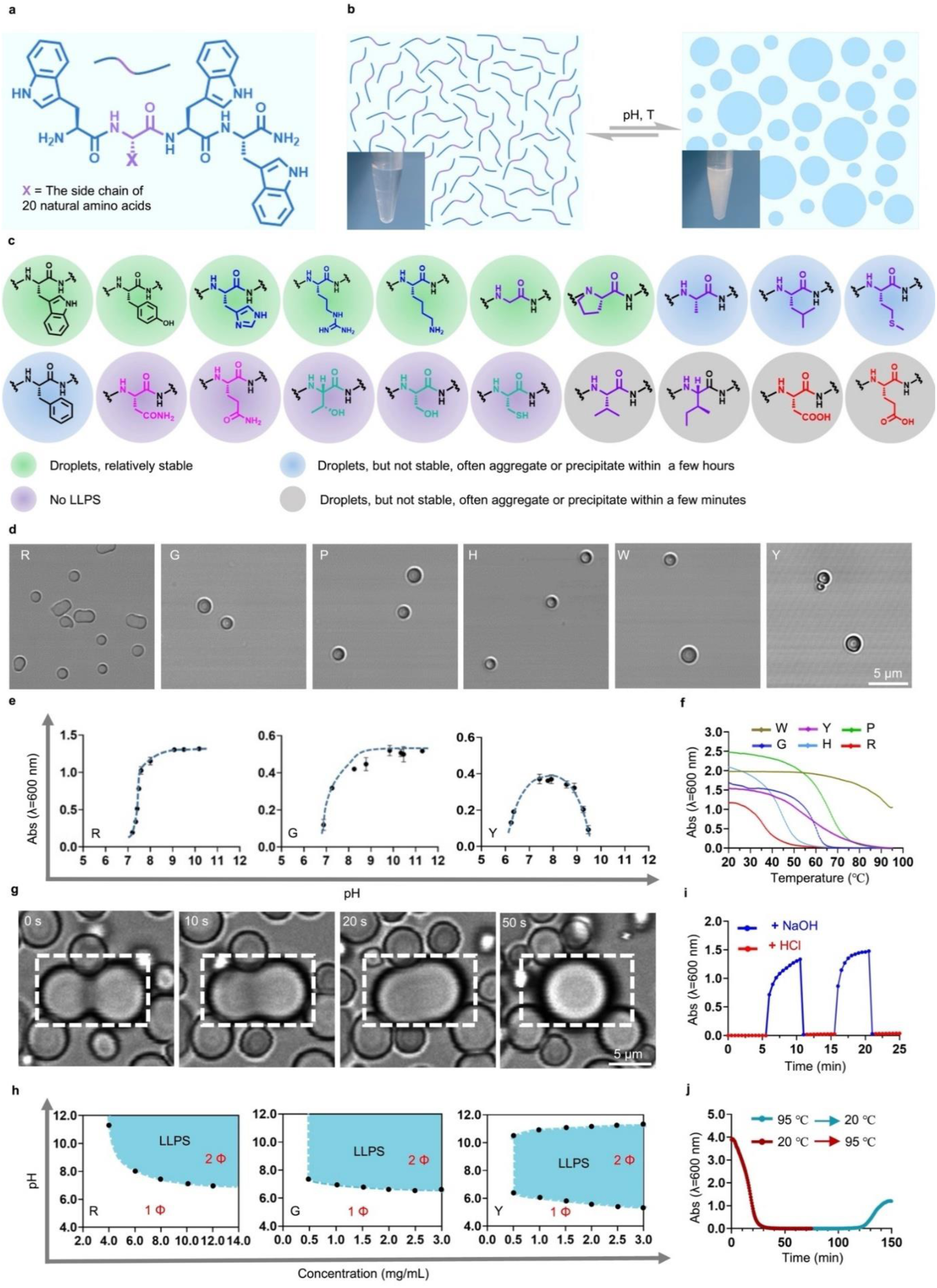
Liquid-liquid phase separation of tetra-peptides. **a**, Chemical structure of tetrapeptide WXWW. **b**, Schematic illustration of liquid-liquid phase separation of tetra-peptide WXWW. **c**, The summary of phase transition behavior of WXWW. **d**, Microscope images of droplets formed by different tetra-peptides (pH= 7.5 ± 0.5). **e**, The turbidity of peptides changes with pH that is measured by spectrophotometer at the absorbance of 600 nm. **f**, Upper critical solution temperature of peptides that measured by spectrophotometer (pH= 7.5 ± 0.5). **g**, The droplets fusion process of WRWW. **h**, pH-concentration phase diagrams of peptides in aqueous solution at room temperature. **i**, pH, and **j**, temperature reversibility curve of WRWW.

The clear solution turned into a turbid emulsion at the optimized pH and temperature. We then measured the pH- and temperature-dependent turbidity by the UV absorption at the wavelength of 600 nm. The results show that their turbidity increased dramatically when pH reached at 7.0, indicating the phase separation was triggered by the deprotonation of the amine group at the N-terminal (Fig. 2e, Fig. S2). The temperature-dependent turbidity demonstrates that the formed droplets all feature an upper critical solution temperature (UCST), and the phase transition temperature of peptides increases as the hydrophobicity or aromaticity of the guest residues increase (Fig. 2f). Time-lapse images acquired from confocal laser scanning microscopy (CLSM) show the fusion of droplets, demonstrating the liquid-like properties of droplets (Fig. 2g, video S1). pH-concentration phase diagrams indicated that the tetra-peptides with the hydrophobic or aromatic guest residue (X = W, Y, P) need a lower pH/concentration to phase separate than the peptide WGWW (Fig. 2h, the right panel, Fig. S4). In contrast, the peptide with the hydrophilic amino acid (X = H, R) needs a higher pH/concentration to form droplets than the peptide WGWW (Fig. 2h, the left panel, Fig. S4). pH and temperature reversibility of the droplets were demonstrated by the measurement of turbidity (Fig. 2i and Fig. 2j). These results demonstrate that tetra-peptides can form liquid-like droplets with adaption by tuning different guest residues.

### The driving force for LLPS of tetra-peptide system

To investigate the driving force that governs droplet formation, we first used Fourier-transform infrared spectroscopy (FTIR) to shed light on the molecular interaction of peptides in LLPS^40, 41^. Before the formation of droplets, the peaks at 1672 cm^-1^ (terminal amide) and the broad bands at 1652 cm^-1^ (internal amide) in the FTIR spectrum indicate no order-organized intermolecular interactions of WRWW (Fig. 3a, blue curve). After the induction of droplets, a significant peak appeared at 1645 cm^-1^, suggesting the increased association of the amide groups between the backbone^42^ (Fig. 3a, red curve). The two peaks at 1611 cm^-1^ and 1586 cm^-1^ belong to the vibration of the aromatic ring shifted to 1606 cm^-1^ and 1580 cm^-1^, respectively, probably because of the π-π stacking of the aromatic rings.

**Fig. 3.**
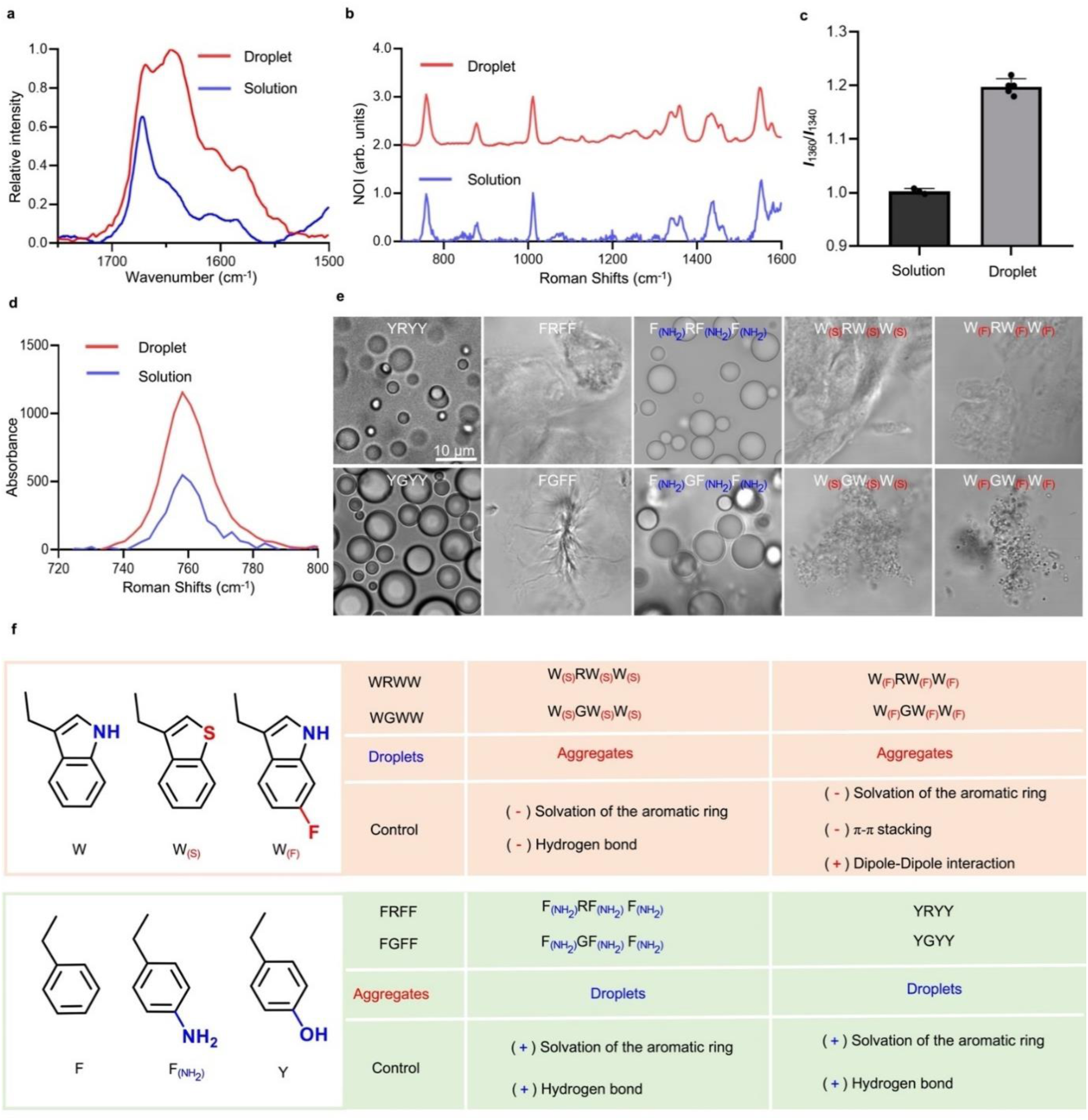
The mechanism underlying liquid-liquid phase separation. **a**, The IR spectra of WRWW in solution and droplets state. **b**, The Raman spectra of WRWW in solution and droplets state. **c**, Hydrophobic interaction in solution and droplet state of WRWW measured by ***I***_1360_/***I***_1340_. **d**, The partially enlarged Raman spectra of WRWW in solution and droplets state. **e**, The bright field images of the microstructure formed by different tetra-peptides. **f**, The relationship between the chemical structure and their microstructure.

We next analyzed the effect of peptide side chains by Raman spectroscopy (Fig. 3b, Fig. S5). The peak around 877 cm^-1^ means that the NH group on the indole ring is involved in a medium-hydrogen bond both in solution and droplet state. For the WYWW, the intensity ratio of the doublet at 850 cm^-1^ and 830 cm^-1^ is about 1.3 (Fig. S6a), indicating moderate hydrogen bonding by the hydroxyl group of tyrosine^43, 44^. The measured Roman shift of 1010 cm^-1^ suggests weak Van der Waals interactions in solution and droplet states. The values of the intensity ratio *I*_(1360)_/*I*_(1340)_ dramatically increased when the droplets formed, indicating stronger hydrophobicity within the droplets than in the solution state^36, 37^(Fig. 3c, Fig. S6b). From the spectra of WRWW, no significant changes near the doublet centered at ∼755 and ∼764 cm^-1^, indicating no cation-π interactions involved with the Trp^38^(Fig. 3d). To further confirm this point, we use the organic base triethylamine (TEA) instead of the sodium hydroxide solution to adjust the pH, excluding the possible cation-π interactions between Na^+^ and Trp. All these peptides can still undergo LLPS (Fig. S2b), further demonstrating that cation-π interactions are unnecessary for the tetra-peptide system.

The above results suggested that the increased hydrophobic interaction, π-π stacking, and the hydrogen bond drive the LLPS of the tetra-peptides cooperatively. To further verify this conclusion and expand the tetra-peptide library for LLPS, we next synthesized the peptides YRYY and FRFF. The results show that peptides YRYY can undergo LLPS, but FRFF forms the aggregates, implying that the hydrophilic group on the aromatic ring is important for LLPS (Fig. 3e, Fig. 3f). To verify the contribution of hydrophilic group on the aromatic ring, we synthesized the peptide W_(S)_RW_(S)_W_(S)_, the NH group on the indole ring was substituted by sulfur (S), which could decrease the hydrogen bonding and water solvation of the peptide. Another peptide F_(NH2)_RF_(NH2)_F_(NH2)_ with amine groups that can provide extra hydrogen bonding and facilitate the solvation of the aromatic ring. As a result, the peptide W_(S)_RW_(S)_W_(S)_ forms aggregates, F_(NH2)_RF_(NH2)_F_(NH2)_ forms droplets, demonstrating that the hydrogen bonding sites on the aromatic ring are essential for the formation of the droplets.

Moreover, the fluorine substituted peptide W_(F)_RW_(F)_W_(F)_ also forms the aggregates, possibly because the π-π stacking of the aromatic rings was disrupted by the increased dipole-dipole interaction or the decreased solvation of the aromatic ring^45, 46^. In parallel, we also synthesized the peptides YGYY, FGFF, F_(NH2)_GF_(NH2)_F_(NH2)_, W_(S)_GW_(S)_W_(S)_, W_(S)_GW_(S)_W_(S)_. They all show the same phase behavior as peptides with R as guest amino acid (Fig. 3e), further demonstrating the great contribution of hydrophilic groups on the aromatic ring for LLPS.

Above all, the interactions among peptide-peptide and peptide-water contribute to the LLPS synergistically. The peptide-water interaction plays a dominant role in the acidic solution state, which leads to better dissolution. With the pH increase, the amine group of the N-terminal began to deprotonate which caused the hydrophobicity of the peptide to increase, following the clustering of the peptide. The increased association of the backbones and side chains work cooperatively to stabilize the droplets. The aromatic rings without the hydrophilic group could hardly interact with the aqueous solvent, the peptide-peptide interaction could not be balanced by peptide-water interaction, resulting in the formation of aggregates rather than droplets. Other factors that disrupt the π-π stacking or increase the hydrophobicity could also cause the formation of the aggregates (Fig. 3f).

### The droplets with core-shell structure

We measured the property of recruiting guest molecules (fluorophores including coumarin 6, SBD, rhodamine 6G, rhodamine B, and BODIPY) from the surrounding milieu and quantified the encapsulation efficiency by fluorescence spectroscopy of the supernatant (Fig. S7). The estimated partition efficiency is about 90%. Specifically, about 99% of the added rhodamine B is encapsulated in the droplets formed by WRWW (Fig. S8). Further observation by CLSM revealed that the fluorescent dye has different distributions within the droplet when added before or after the droplet formation (Fig. 4a). When the addition of fluorophore before the droplet formation, the fluorescent molecules were mainly located at the inner core of the droplets (Fig. 4b top panel, Fig. S9). Meanwhile, with the addition of fluorophore after the droplet formation, we found the droplets exhibit a vesicular-like structure (Fig. 4b bottom panel). These results together implied that the droplets could have a core-shell structure, which could spatially control the distribution of guest molecules (Fig.S10).

**Fig 4.**
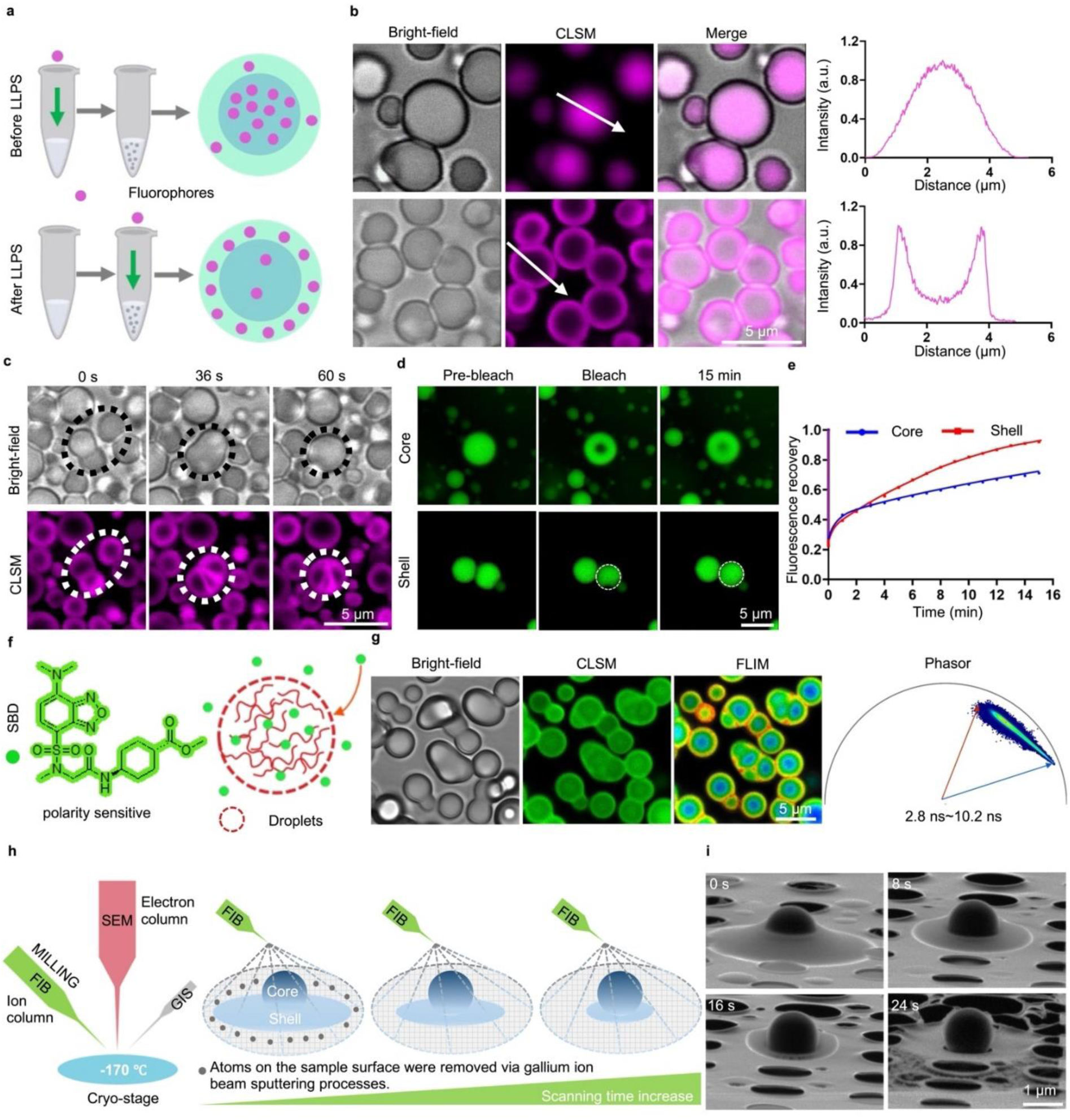
The core-shell structure of the droplets. **a**, Schematic illustration of the fluorophores added before and after the droplet formation. **b**, The CLSM images of the rhodamine B added before and after the droplet formation. **c**, Time-dependent CLSM images show the changes of the inner core during the fusion process of the outer shell. **d**, **e**, Fluorescence recovery after photobleaching (FRAP) experiment. **f**, Chemical structure of polarity sensitive molecule SBD and the schematic illustration of it enter into the droplets. **g**, DIC, CLSM, FLIM, and Phasor images of the droplets after adding SBD. **h,** Schematic illustration of the Cryo-FIB/SEM hardware component and the changes of the droplets under the gallium ion beam sputtering. **i,** Time-dependent Cryo-FIB images of the droplets under the gallium ion beam sputtering.

Since the bright field images could not recognize the core-shell structure, we tried to examine the fusion process by CLSM. The timelapse images (video S2) show that the inner core of the droplets has been squeezed to deform rather than fuse during the fusion processes of the outer shell, suggesting that the inner cores of droplets are not solid materials but gel-like matrix (Fig. 4c). We next performed the fluorescence recovery after the photobleaching (FRAP) experiment at the center and periphery of the droplets. The fluorescence recovery rate of the inner core is much lower than that of the shell, suggesting that fluorescent molecules experience stronger molecular interaction within the inner core (Fig. 4d, e). To investigate the difference in the microenvironment of the inner and shell of the droplets, we used fluorescence lifetime imaging microscope (FLIM) experiments by using sulfonamide benzoxadiazole (SBD), which is a polarity-sensitive dye that could report the average microenvironmental polarity in close proximity to the fluorophores by exhibiting distinctive fluorescence lifetimes^47^ (Fig. 4f). The results show that the lifetime of SBD gradually decreases from the outer shell to the inner core, demonstrating that the droplets are inhomogeneous (Fig. 4g, video S3). To exclude the possible interpretation of lifetime that caused by the indole ring from the Trp which is also polarity sensitive, or the trapped water of the inner core, we used the Cryo-EM, Cryo-SEM combined with Cryo-FIB to observe the core-shell structure directly. The results show that the spherical core is surrounded by the collapsed outer shell (Fig. S11). The time-dependent Cryo-FIB images indicate that the outer part of droplets quickly disappeared because the atom of the sample can be continuously removed via the ion beam sputtering process. Meanwhile, the inner core is relatively stable when simultaneously exposed to the same gallium ion beam (Fig. 4h, 4i, Fig. S12, Fig. S13). This result indicates that the molecules are packed more tightly or interact more strongly with molecules in the inner core than in the outer shell.

### The strong π-π interaction defines the core-shell structure

Stress granules have a similar core-shell structure with a stable inner core surrounded by a liquid-like shell^4, 48, 49^. During the liquid-liquid phase separation, the reversible and dynamic fibers formed in the inner core, which can regulate the fluidity and mobility of the inner core, eventually causing the formation of hydrogel-like cores or reversible amyloid cores^50, 51^. Although the tetra-peptide usually tends to self-assemble into hydrogels with fiber structure rather than liquid-liquid phase separation^52^, we hypothesized that the LLPS of peptides in our system may still reserve the propensity for fiber formation. Changing the amino acid composition from W to other amino acids, such as Y, can form homogeneous droplets instead. A very slight change in the crosslinks at the molecular level can dramatically strengthen or weaken the percolation or self-association of the condensed phase^32, 53, 54^. Therefore, the stronger interaction between W may account for forming gel-like cores.

To verify our hypothesis and figure out what kinds of driving forces cause the core-shell structure, we synthesized the peptide W_(OH)_RW_(OH)_W_(OH)_ and W_(DHT)_RW_(DHT)_W_(DHT)_. We also changed the amino acid from W to other amino acids such as Y (Fig. 5a). The peptide W_(OH)_RW_(OH)_W_(OH)_ increased 3 hydrogen bonding sites on each molecule compared with the peptide WRWW, which does not substantially affect the core-shell structure (Fig. 5b left panel). We aborted the double bond on the indole ring to form W_(DHT)_RW_(DHT)_W_(DHT)_, which is no longer conjugate to the benzene ring, thus reducing the aromatic interaction compared with WRWW. The CLSM and fast FLIM results all show that W_(DHT)_RW_(DHT)_W_(DHT)_ forms homogeneous droplets (Fig. 5b middle panel). The amino acid Y weakens the crosslink of molecules within the condensed phase during the liquid-liquid phase separation because it shows weakened π-π interactions than W. The results show that YRYY formed homogeneous droplets (Fig. 5b right panel). Moreover, the parallel group WGWW, W_(OH)_GW_(OH)_W_(OH)_, W_(DHT)_GW_(DHT)_W_(DHT)_, YGYY show similar results with the R group (Fig. S14). These results suggested that the strong π-π interaction is the main driving force for the formation of dynamic fibers and eventually defines the core-shell structure of the droplet, which is consistent with the rules of peptide self-assembly that aromatic-aromatic interactions are beneficial to the formation of fibers or hydrogels.

**Fig. 5.**
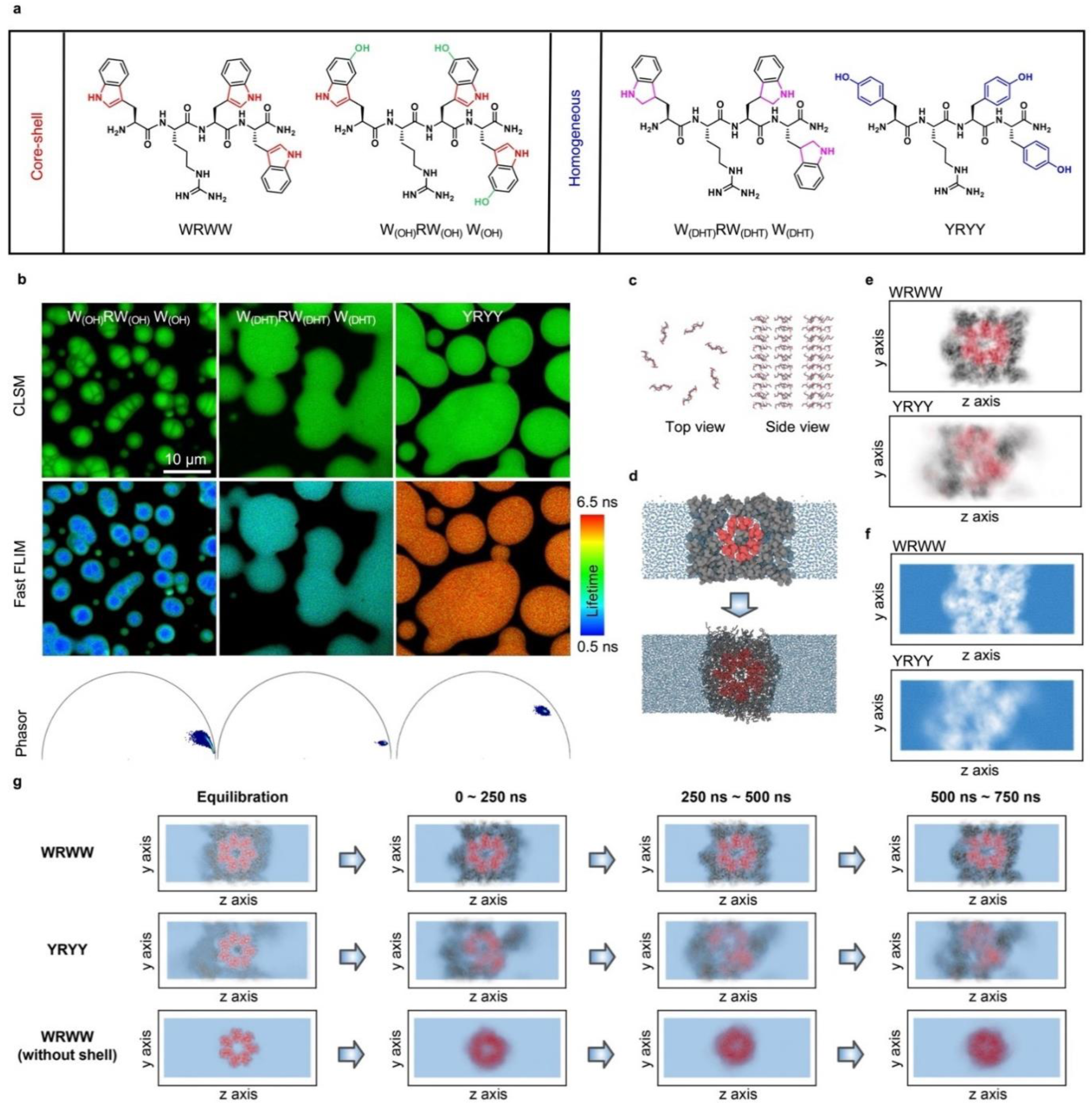
The mechanism underlying the formation of the core-shell structure. **a**, Chemical structure of the peptides WRWW, W_(OH)_RW_(OH)_W_(OH)_, W_(DHT)_RW_(DHT)_W_(DHT)_, YRYY. **b**, The confocal and fast FLIM images of the peptides W_(OH)_RW_(OH)_W_(OH)_, W_(DHT)_RW_(DHT)_W_(DHT)_, YRYY. **c,** Backbone structure of the fiber-like core. d, Multiscale simulation scheme. The initial structure of all-atom simulation (bottom) was prepared using MARTINI simulation (top). The color coding is consistent across all panels: core peptides (red), shell peptides (black), and water molecules (blue). **e,** Average peptide density profile for the last 250 ns of the simulation in a basic environment. **f,** Average water density profile for the last 250 ns of the simulation in a basic environment. **g,** Average density profile of WRWW, YRYY and WRWW (without shell) across different simulation time windows.

To further support our hypothesis, a multiscale simulation approach was adopted to efficiently and accurately simulate tetra-peptide systems, as illustrated in Fig. 5d. Based on the fiber-like core hypothesis, we constructed the initial configuration of the tetrapeptide systems as a central core with several anti-parallel tetrapeptide branches (Fig. 5c), surrounded by shell peptides (Fig. 5d, Fig. S15). The pKa of uncapped N-termini in peptides is approximately 8.0^55^, implying that under acidic conditions, the N-termini of tetrapeptides are protonated, while at basic conditions, the N-termini of tetrapeptides are deprotonated. We simulated both acidic and basic environments for the WRWW and YRYY systems, resulting in four distinct simulations. The multiscale simulation started with a 1 μs long coarse-grained simulation using the MARTINI 3 model^56^, followed by a more detailed 750 ns all-atom simulation using the CHARMM36m force field^57^. To analyze the distribution of tetrapeptide molecules in the simulation box, we computed the average probability density of various atoms across the *yz*-plane throughout the all-atom simulation process.

The simulation results were in agreement with experimental observations. In an acidic environment, both WRWW and YRYY systems transitioned to a single, homogeneous dilute phase, with the condensed phase absent. In contrast, the condensed phase persisted in both systems in a basic environment (Fig. S16). Notably, in a basic environment, the WRWW fiber core maintained its original structure in the droplet, while the initial fiber core dissipated in the YRYY droplet (Fig. 5e). The consistency between simulation and experimental results supports the validity of the fiber-core and liquid-shell model. Additional simulations conducted on WGWW and YGYY yielded analogous results, which are reported in the Supplementary Information (Fig. S16, Fig. S17, and Fig. S18).

In addition to simulations of the fiber-like core within a liquid shell, we conducted a simulation for a bare WRWW core without a surrounding liquid shell to investigate the rationale behind the formation of the core-shell microstructure. Comparison of the fiber density distributions of the WRWW core during the simulation, both with and without the liquid shell, revealed that the absence of a liquid shell allowed more solvent water molecules to penetrate between peptides, leading to the gradual dissolution of the fiber structure (Fig. 5g). This observation provides insight into the existence of the core-shell structure in WRWW condensates: water molecules, interacting at the exterior of the droplets, exert a significant influence on peptide-peptide interactions. These interactions disrupt the ordered fiber structure, inducing a transition towards a disordered liquid state at the exterior of droplets. The equilibrium between peptide-peptide and peptide-water interactions results in a disordered liquid shell encapsulating the solid fiber core.

### Dynamic control the LLPS and the transformation between multiphasic and homogeneous droplets

We have shown that the absence of a hydrophilic group in the aromatic ring often causes the tetrapeptides unable to LLPS because the strong peptide-peptide interaction cannot be balanced by the peptide-water interaction, whereas the introduction of a hydrophilic group could rescue them. The tetra-peptide with stonger peptide-water interaction is also unable to form droplets and remains in a solution state even at a very high concentration. To rescue this type of tetra-peptide, increasing the peptide-peptide interaction by combining two tetra-peptide molecules into one octa-peptide through a disulfide bond could be a quite feasible method, which could dramatically increase the valence of each monomer peptide and decrease the minimal concentration needed for LLPS^31, 32^. Disulfide bonds formed due to oxidation of the thiol group (SH group) come from cysteine (C) residue, which is extensively present in many types of protein molecules, the formation and cleavage of disulfide bonds has been widely used by living systems to govern the basic biological process. Since disulfide bonds can be formed or broken by oxidation–reduction process by adding many kinds of oxidizing and reducing agents, we use hydrogen peroxide (H_2_O_2_) and dithiothreitol (DTT) to control the LLPS dynamically by controlling the formation and cleavage of disulfide bonds (Fig. 6a). The peptide YCGY has a very good solubility in aqueous solution and can’t undergo LLPS even at 80 mg/mL. However, it can undergo LLPS at a low concentration of 6 mg/mL through the formation of disulfide bonds by oxidation of the thiol group. The droplet emulsion turns into a clear solution again after adding DTT (Fig. 6b). The reversible disulfide bond enables us to dynamically control the LLPS. Besides the redox reaction, we can also control the LLPS by post-translational modification of peptides through phosphorylation (Figure S19). Moreover, the transformation between multiphasic and homogeneous droplets can also be controlled by disulfide bonds (Fig. 6c, Fig. 6d). The peptide WCRY forms homogeneous droplets, which transform into the multiphasic droplets after oxidation by H_2_O_2,_ because of the increased π-π interactions by cross-linking. Interestingly, the homogeneously distributed guest fluorescent molecules were recruited into the newly formed inner core, which is very similar to the cellular multiphasic condensates to recruit different biomacromolecules into different layers for fulfilling a variety of functions.

**Fig. 6.**
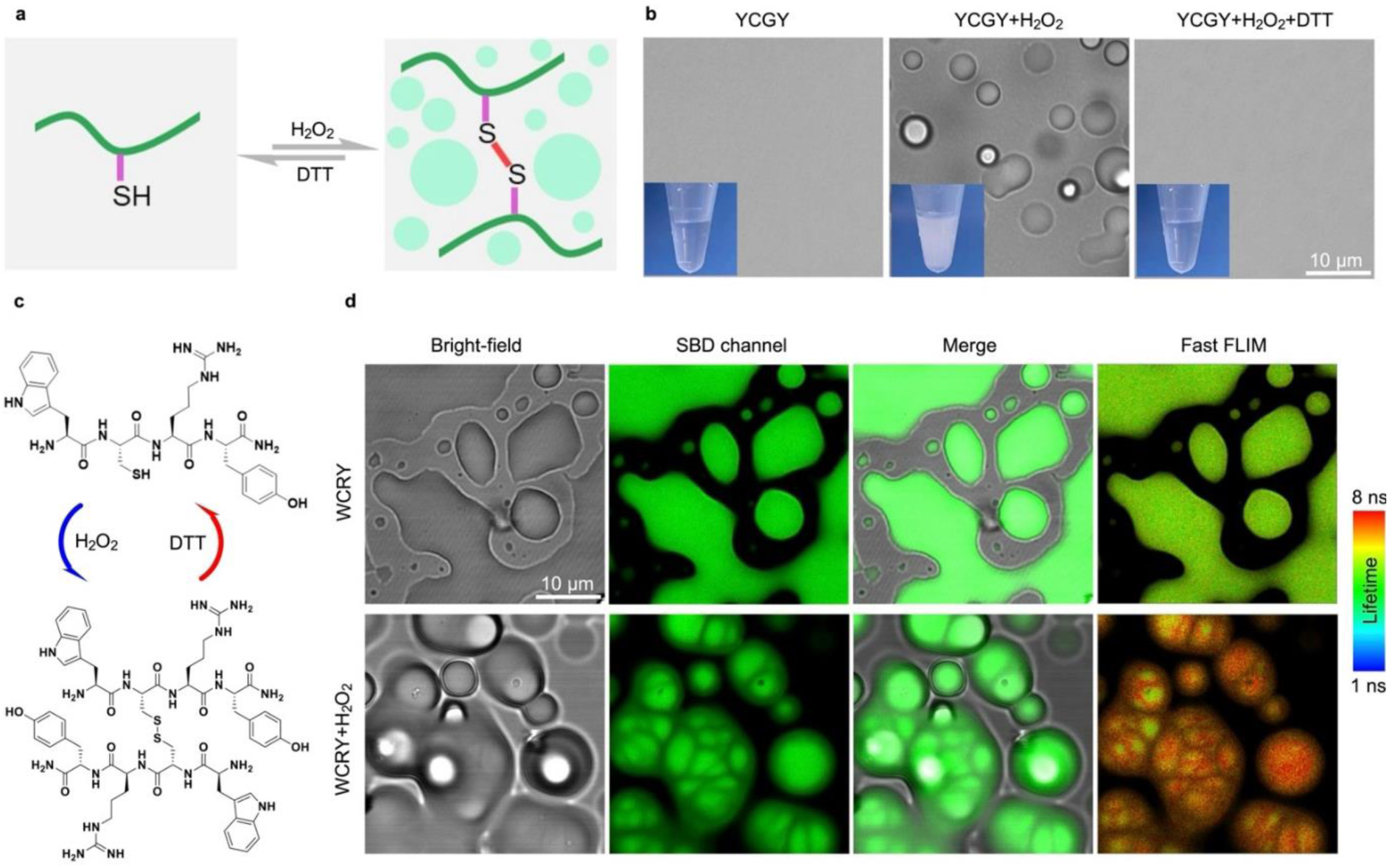
The dynamic and reversible control of LLPS and core-shell structure of the droplets. **a**, Schematic illustration of redox reaction control the LLPS. **b**, the bright-field images of the peptide solution, oxidized and cross-linked by H_2_O_2_, then reduced by DTT. **c**, Schematic illustration of the homogeneous droplets and droplets with core-shell structure can be tuned by redox reaction. **d**, the confocal and fast FLIM images of the homogeneous droplets and the core-shell structure of the droplets formed by the oxidation of H_2_O_2_.

## Conclusions

In summary, this work describes symmetrical core-shell structural droplets through the LLPS by chemical programming. The single component of tetra-peptide can synchronously assemble into a liquid spherical core-shell structure under the external stimuli of pH or temperature. Hydrophobic interaction, π-π stacking, and hydrogen bonds synergistically contribute to droplet formation. The hydrophilic group in the aromatic ring is essential for LLPS because removing it often leads to the formation of aggregates. Moreover, the results suggested that the amino acid W is critical for forming core-shell structure because of its stronger π-π interaction compared with other amino acids. By encoding the amino acid composition of the tetra-peptides, we can design and synthesize not only homogeneous droplets but also droplets with core-shell structures. Finally, we show that homogeneous droplets can transform into the core-shell structure under external stimuli. At the same time, the homogeneously distributed guest molecules were recruited into the inner core, which allows us to control the distribution of guest molecules spatiotemporally. This work facilitates the rational design of synthetic core-shell droplets with various applications on demand and opens a door for synthesizing multiphasic droplets with a single component.

## Methods

### Material sources

Fmoc-amino acids, HBTU (O-(Benzotriazol-1-yl)-N, N, N’, N’-tetramethyluronium hexafluorophosphate) and Rink Amide resin were obtained from GL Biochem (Shanghai, China). Rhodanmin B, rodanmin 6G, trifluoroacetic acid (TFA), triethylamine (TEA), N, N-diisopropylethylamine (DIPEA), NaOH, and triisopropylsilane (TIS) were purchased from Aladdin (Shanghai, China). Coumarin 6 and fluorescein were purchased from Macklin (Shanghai, China). N, N’-dimethyl formamide (DMF) was obtained from J&K Scientific (Beijing, China). All other reagents were purchased from commercial sources and used without further purification.

### Peptide synthesis

All peptides were synthesized by Fmoc based solid-phase peptide synthesis (SPPS) (Fig. S1). Briefly, after the rink amide resin was swelling in DMF, the first amino acid with side-chain protected in DMF was loaded to the resin by coupling reagent HBTU and DIPEA for 2h. After washing 5 times by DMF, we used deprotection agent (20% piperidine in DMF) to remove the Fmoc group. Then the growth of the peptide chain follows the established Fmoc SPPS protocol. The prodcut of the peptide was cleaved from the resin by using cleavage reagent (TFA: TIS: H_2_O = 95%: 2.5%: 2.5) for 2 hours. After the solvent was removed through the rotary evaporation, ice-cold diethyl ether was added to the above residue. The resulting precipitate was centrifuged at 5000 g for 3 minutes at 4 °C. The supernatant was removed, the resulting crude peptide was further purified by High Performance Liquid Chromatography (HPLC).

### The preparation of peptide droplet

The lyophilized tetra-peptide powder was dissolved in Milli-Q water and then adjusting the pH gradually to 7.5 by adding 0.5 M NaOH (aq.). The resulting milky emulsion was heat at 60 ℃ for 5 minutes and cooled to room temperature. Then the milky emulsion was examined by bright-field microscopy (Leica STELLARIS 8 FALCON confocal microscope). The final concentration (Table S1) of tetra-peptides determined by their hydropathy and aromatic property.

### Turbidity measurement

We measured the turbidity of all the peptide solution by spectrophotometer at the wavelength of 600 nm. All the measurments were carried out for three independent times at room temperature. The 0.1 M, 0.5 M, 1.0 M NaOH was chosen depending on the concentration of different peptides and the added volume did not exceed 10% of the original volume. The same volume of Milli-Q water was used as control group. The pH-dependent turbidity measurements were performed on a Microplate reader (Thermo Scientific Varioskan LUX). Other turbidity were measured by Agilent carry3500. The pH was measured by pH-meter (METTLER TOLEDO FE28 -Micro).

### pH-concentration phase diagrams

The pH-Concentration phase diagrams was measured in the following steps: we set 6 different concentrations for each peptide. For each specific concentration, 160 μL peptide solution was prepared, then gradually increase the pH by adding NaOH (aq.) to the peptide solution and monitor the turbidity changes by spectrophotometer at the wavelength of 600 nm (Agilent carry3500). When a turbidity appeared, measure the pH by pH-meter (METTLER TOLEDO FE28 -Micro).

### Fourier transform infrared spectroscopy of peptide droplets or the solution

The lyophilized tetra-peptide powder was dissolved in deuterium oxide (D_2_O) and then adjusting pH to 7.5 by 0.5 M sodium deuteroxide (NaOD). Using a pipettee to drop the solution or droplets of the peptide on a diamond single reflection attenuated total reflectance (ATR) module and examine their spectra by the FTIR micro spectrometer (ThermoFisher Nicolet iS50) with averaging 16 scans, spectral resolution of 4 cm^-1^.

### Raman spectroscopy of peptide solution and the droplets

10 μL of the samples was settled on the glass slide with precoated sigmacote. For the peptide solution, average Roman spectra was obtained by VIS-NIR Confocal Raman Microscope System (WITec alpha 300R). For the peptide droplets, we used a 50×magnification objective lens to focuse the droplets at the bottom layer. Then we obtained their Raman spectra by using a 532 nm laser excitation with 10 mW laser power with a 10 s integration time and 10 times accumulation. For each sample, 5 different droplets was measured for statistic analysis.

### Encapsulation efficiency of guests by droplets

After the formation of droplets, 1 μL solution of fluorophore was added to100 μL droplets emulsion and the mixture was incubated for 1 h at rt., then the samples were centrifuged at 3,000 × g for 20 min, then 20 μL of supernatant was diluted to 100 μL. Rodanmin 6G, Rodanmin B were diluted with water, Coumarin 6, Resorufin Sodium Salt, SBD, Red BODIPY were diluted with DMSO. The concentration of the fluorophores at the supernatant was measured via fluorescence. Efficiency of encapsulation (%EE) was calculated using %EE = C_Total_ - C_supernatant_ / C_Total_. We tested the encapsulation of WYWW at pH 9.0 because of the droplets formed by WYWW tend to aggregates at pH 7.5 when the fluorophores were added, but relatively stable at pH 9.0.

### Fluorescence recovery after photobleaching (FRAP)

After the formation of droplets, 0.1 μL solution of fluorescein (1 mM, DMSO) was added. About 10 μL droplets emulsion was settled on the glass slide with precoated sigmacote and left for 16 hours before the FRAP experiment to allow the small droplets to fuse into the big one. The photobleach and fluorescence recovery was recorded by a Leica STELLARIS 5 FALCON confocal microscope at the excitation wavelength of 488 nm.

### Fast fluorescence lifetime imaging microscopy (FLIM) of droplets

A drop of phase-separated solutions was settled on the glass coverslip for imaging. The FLIM images were acquired by a Leica STELLARIS 8 FALCON confocal microscope equipped with a pulsed white laser (WLL) using an X63/1.40 oil immersion objective. The SBD was excited by a 448 nm laser line with a frequency of 10 MHz, respectively. The fluorescent lifetime fitting and image analysis were performed in LAS X and Fiji.

### Samples preparation for Cryo-EM, Cryo-SEM, Cryo-FIB

About 3 μl droplets emulsion were applied onto glow-discharged holey carbon grids (Quantifoil Cu R1.2/1.3, 300 mesh), blotted with a Vitrobot Marker IV (Thermo Fisher Scientific) for 3 s under 100% humidity at 4 °C, and subjected to plunge freezing into liquid ethane.

### Cryo-FIB milling

Vitrified samples were further processed by cryo-FIB milling. A dual-beam microscope FIB/SEM Aquilos 2 (Thermo Fisher Scientific) equipped with a cryo-transfer system (Thermo Fisher Scientific) and rotatable cryo-stage cooled at −191 °C by an open nitrogen circuit was used to carry out the thinning. Prior to the milling, the grids were mounted on the shuttle and transferred onto the cryo-stage, followed by the coating with an organometallic Platinum layer using the GIS system (Thermo Fisher Scientific) for 5–6 s. Then, droplets positioned approximately in the centres of grid squares were selected for thinning. Thinning was carried out under the condition of voltage 30kV and current 10pA,until the material around the ball is burned off.

### Redox controlled droplets formation through LLPS

The lyophilized tetra-peptide powder YCGY was dissolved in Milli-Q water, then the pH of resulting homogeneous solution (200 mL, 10 mg/mL) was adjusted to 7.5 by using 0.5 M NaOH. Then 5 uL was taken out and examined by CLSM. Then 2 μL H_2_O_2_ (1M) was added to the remain residual solution. After incubation at room temperature for 5 mins, 5 μL was taken out for obsevation by CLSM. Then 2.5 DTT (1M) was added to the residual liquid and incubated for 5 mins at room temperature before abserving by CLSM.

### Redox controlled the droplet transformation between homogeneous and core-shell structures

The lyophilized tetra-peptide powder WCRY was dissolved in Milli-Q water and adjusted the pH to 7.5 using 1 M NaOH. After the droplets formation, 0.1 μL SBD (1 mM, DMSO) was added to 100 μL (30 mg/mL) droplets emulsion and incubation at room temperature for 5 mins. Then 5 μL was taken out for the obsevation by CLSM and fast FLIM imaging. Then 1 μL H_2_O_2_ (1M) was added to the remain residual solution and incubation at room temperature for 5 mins before confocal and fast FLIM iamging.

### Multiscale Simulations for Tetrapeptide Droplets

Full details of multiscale simulations for tetra-peptide droplets are given in the Supplementary Information.

## Acknowledgements

This project was supported by the National Natural Science Foundation of China (82272145) and the Foundation of Westlake University. We thank the Instrumentation and Service Center for Molecular Sciences, Instrumentation and Service Center for Physical Sciences, General Equipment Core Facility, Biomedical Research Core Facilities, and Laboratory Animal Resource Center at Westlake University for the assistance with measurements. We acknowledge X. J. Wang for help with the Cryo-FIB experiments and thank Z. Chen for help with the Roman experiments.

## Author contributions

H.M.W. conceptually designed the strategy for this study, provided intellectual input, supervised the studies, and revised the manuscript. Lai.C. Z. designed the study and perform most of the experiments. Long.C.Z and X. Z. performed the FLIM experiments. C.W. and B. Z. did MD simulations. T.Y.X. and J.W. synthetized some peptides and perfomed CLSM experiments. H.M.W. and Lai. C.Z. wrote the manuscript with contributions from all authors.

## Data and materials availability

All the data that support the findings are available within the main text and the supplementary materials.

