## Supplementary information for "Synchronous assembly of peptide anisosome"

**Description of video**

Supplementary Video 1. Time-dependent bright field imaging of WRWW (18 mg/mL, pH 7.5 ± 0.5) shows the fusion process of the droplets. Imaging was conducted with 1-frame-per-5-second rate, related to figure 2 g.

Supplementary Video 2. Time-dependent CLSM imaging of WRWW (18 mg/mL, pH 7.5 ± 0.5) droplets with Rho B (1 μM) added after droplets formation, which shows the deformation of inner core during the fusion processes of the outer shell. Imaging was conducted with 1-frame-per-2-second rate, related to figure 4 c.

Supplementary Video 3. Three-dimensional FLIM images of WRWW (12 mg/mL, pH 7.5 ± 0.5) droplets with SBD (1 μM). The sectional view shows the heterogeneous core localizes the center of the droplets, related to figure 4 g.

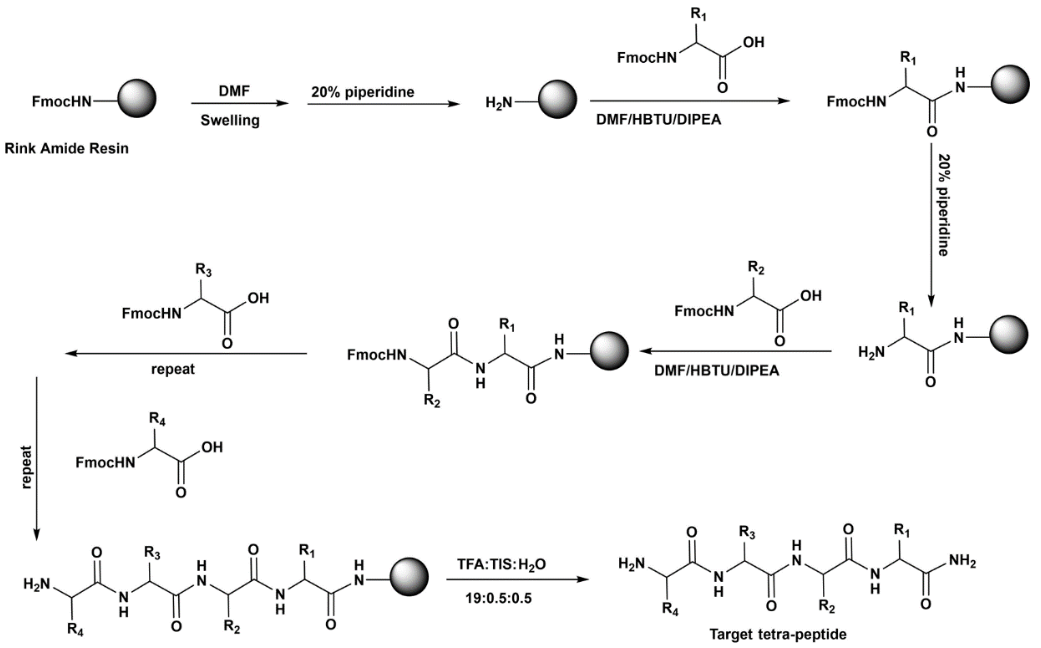


**Supplementary Fig. 1:** The synthetic routes for solid phase peptide synthesis of tetra-peptides.

| Concentration (mg/mL) | Peptide |
|---|---|
| 0.5 | $W_{(S)}GW_{(S)}W_{(S)}$, $W_{(F)}GW_{(F)}W_{(F)}$ |
| 1 | WWWW, WYWW, WFWW, WLWW, WIWW, WVWW, WMWW, WPWW, WAWW, WGWW |
| 2 | WHWW, WCWW, WSWW, WTWW, WNWW, WQWW |
| 4 | WKWW, $W_{(OH)}GW_{(OH)}W_{(OH)}$ |
| 6 | WDWW, WEWW |
| 8 | FGFF, FRFF, $W_{(S)}RW_{(S)}W_{(S)}$, $W_{(F)}RW_{(F)}W_{(F)}$, WGpYW |
| 10 | YCGY |
| 12 | WRWW |
| 15 | $W_{(DHT)}GW_{(DHT)}W_{(DHT)}$ |
| 30 | YGYY, YRYY, WCRY, $W_{(OH)}RW_{(OH)}W_{(OH)}$, $W_{(DHT)}RW_{(DHT)}W_{(DHT)}$ |
| 100 | $F_{(NH_2)}GF(NH_2)F(NH_2)F(NH_2)$ |
| 150 | $F_{(NH_2)}RF(NH_2)$ $F(NH_2)F(NH_2)$ |

**Supplementary Tab. 1:** The final concentration for LLPS of tetra-peptides.

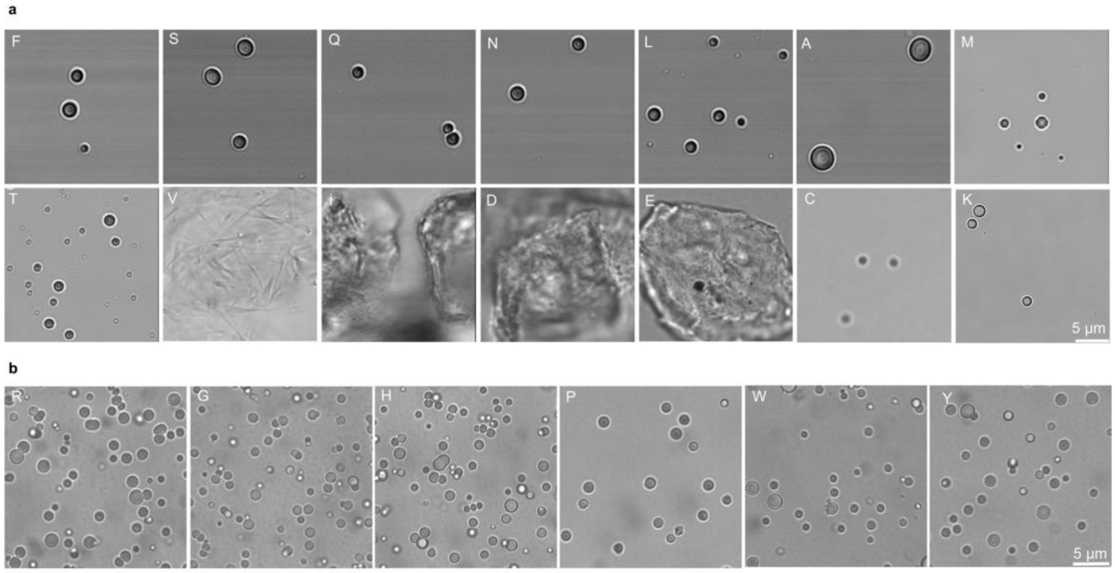


**Supplementary Fig. 2:** The bright field images of WXWW. **a,** The pH was adjusted by 0.5 M NaOH. **b,** The pH was adjusted by 0.5 M triethylamine (TEA). The final pH was 7.5 and final concentration are seen in Table S1.

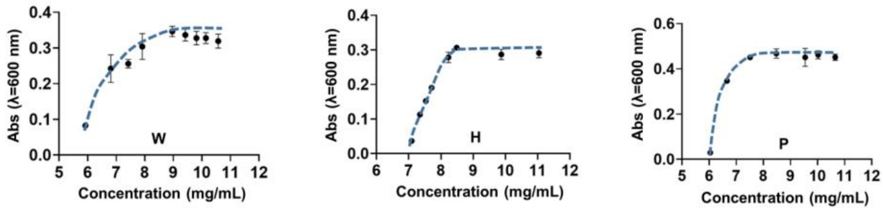


**Supplementary Fig. 3:** The turbidity of peptides changes with pH that measured by spectrophotometer at the wavelength of 600 nm.

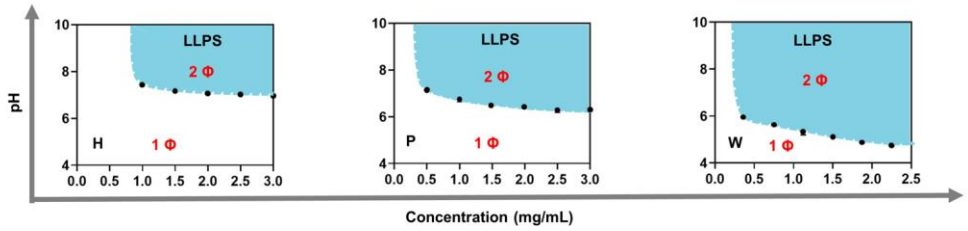


**Supplementary Fig. 4:** pH-concentration phase diagrams of peptides in an aqueous solution at room temperature.

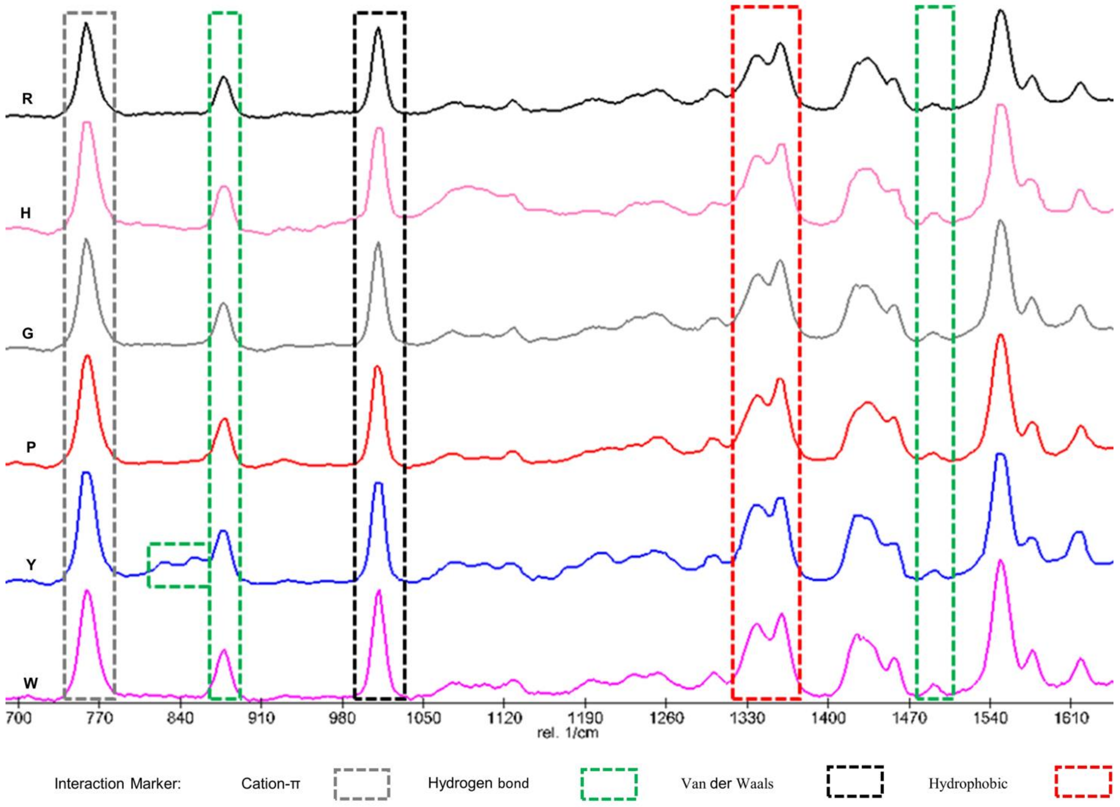


**Supplementary Fig. 5:** The Raman spectra of WXWW in Droplets state.

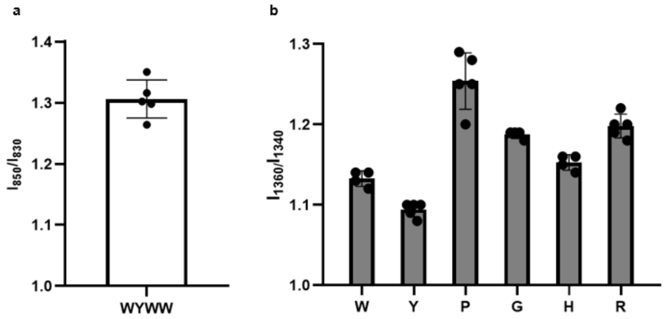


**Supplementary Fig. 6: a,** Hydrophobic interaction in solution and droplets state of WXWW measured by $\boldsymbol{I}_{1360}/\boldsymbol{I}_{1340}$. **b,** The strength of hydrogen bond measured by $\boldsymbol{I}_{850}/\boldsymbol{I}_{830}$ of WYWW.

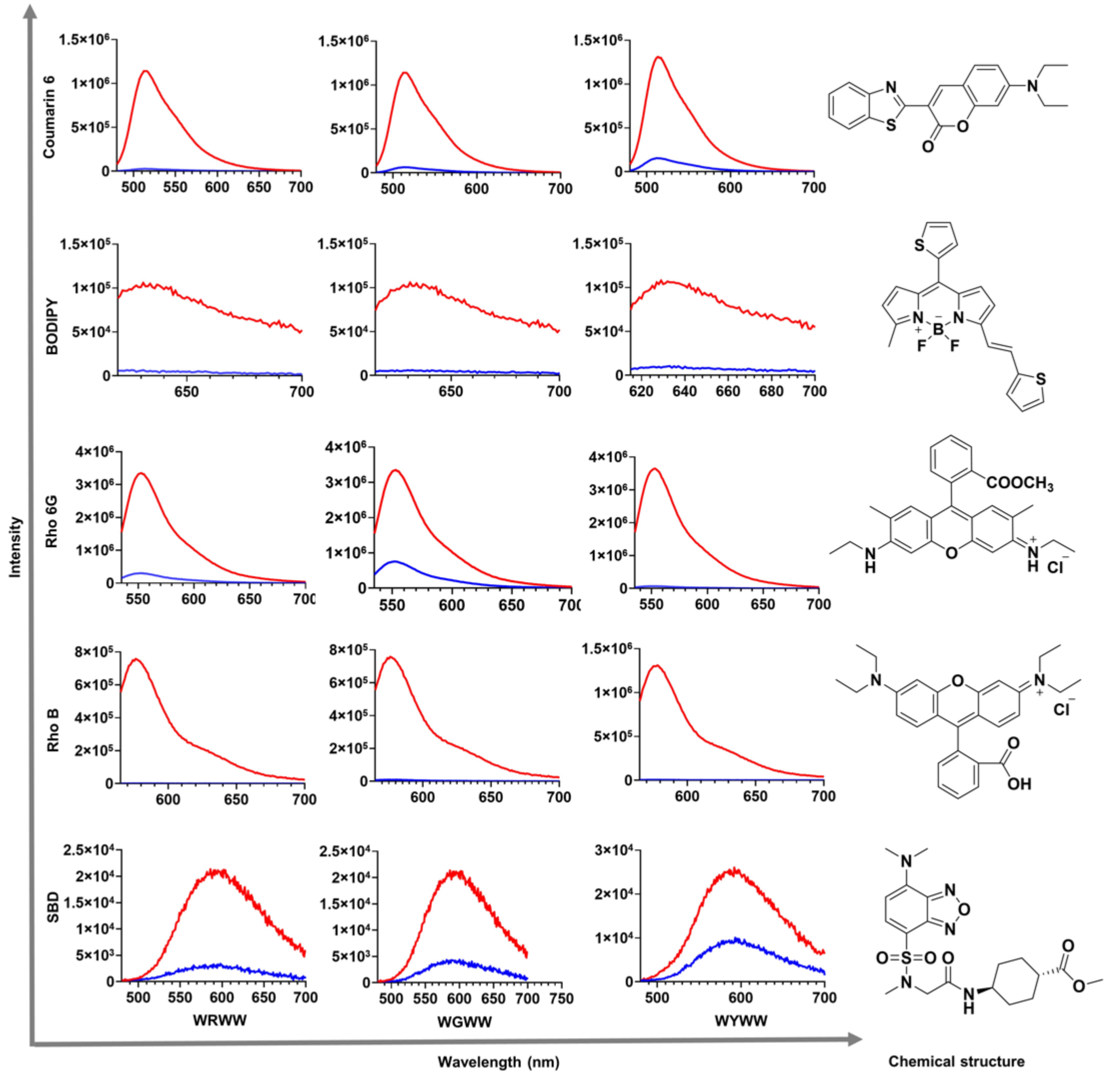


**Supplementary Fig. 7:** The fluorescent spectra of supernatants of buffer（blue curves）and the solutions before phase separation (red curves) containing coumarin 6, BODIPY, rhodamine 6G, rhodamine B, and SBD. Condition: WRWW (12 mg/mL, pH 7.5),

WGWW (1 mg/mL, pH 7.5), WYWW (1mg/mL, pH 9.0), Fluorescent dyes = 10 μM, aqueous solution.

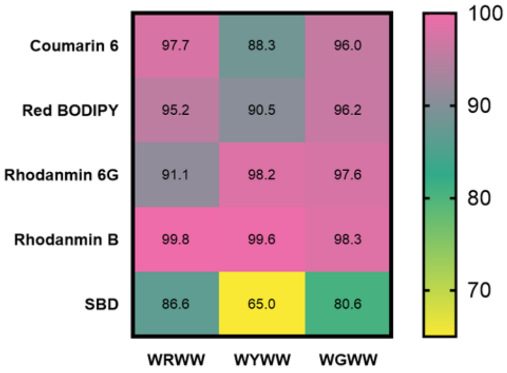


**Supplementary Fig. 8:** The calculated partition efficiency of the fluorescent dyes uptake by the droplets of WRWW, WGWW, WYWW.

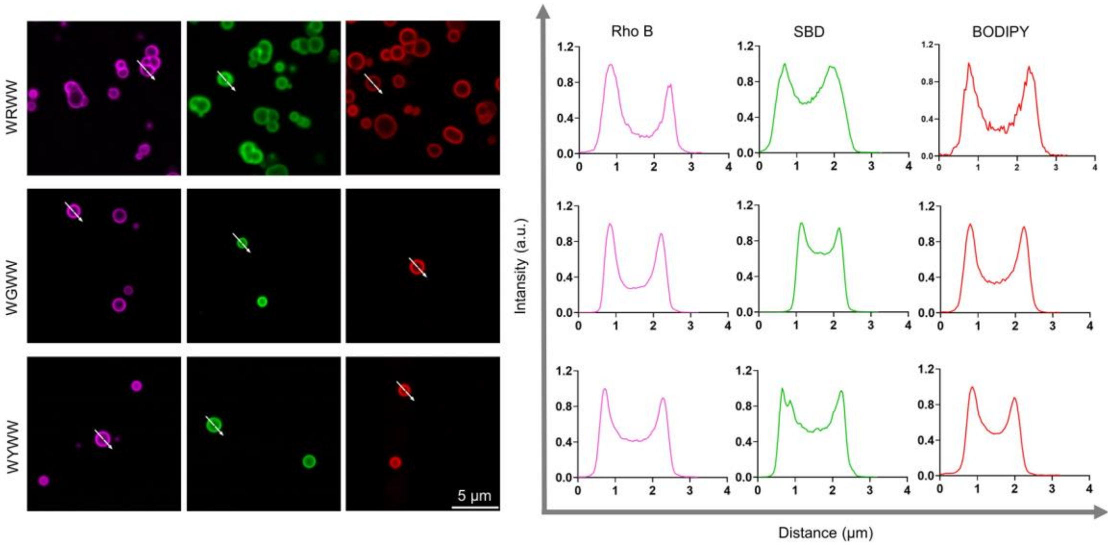


**Supplementary Fig. 9:** The CLSM images of the fluorophores were added to the WGWW, WYWW after the droplets formation and the fluorescence intensity plots at cross-sections.

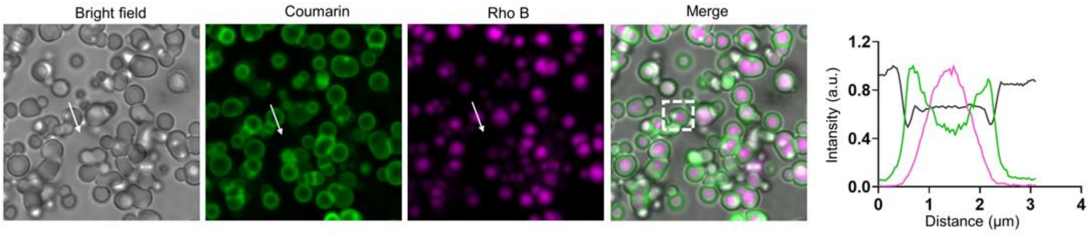


**Supplementary Fig. 10:** Spatiotemporally control the distribution of guest molecules in the droplets. Rhodamine B was added before the droplets formation, Coumarin was added after the droplets formation. WRWW (12 mg/mL, pH=7.5)

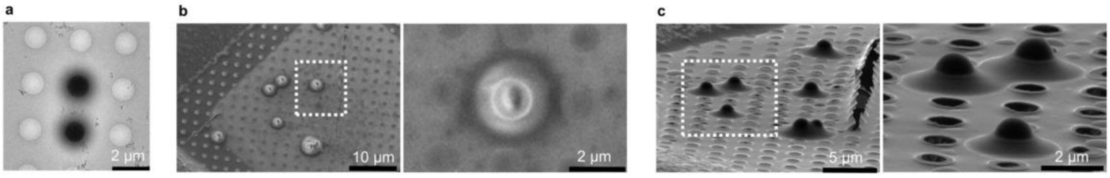


**Supplementary Fig. 11: a, b, c,** Cyro-EM, Cryo-SEM, Cryo-FIB images of the droplets formed by WRWW (12 mg/mL, pH= 9.0)

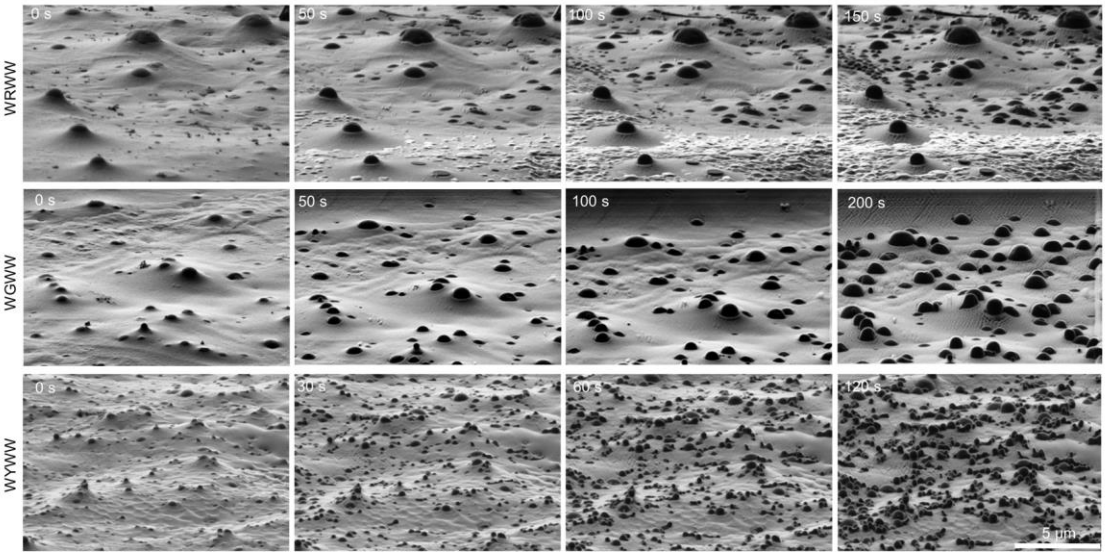


**Supplementary Fig. 12:** Time-dependent Cryo-FIB images of the droplets formed by WRWW, WGWW, WYWW under the gallium ion beam sputtering. (WRWW, 12 mg/mL, pH 7.5, WGWW, 1 mg/mL, pH 7.5, WYWW, 1 mg/mL, pH 9.0)

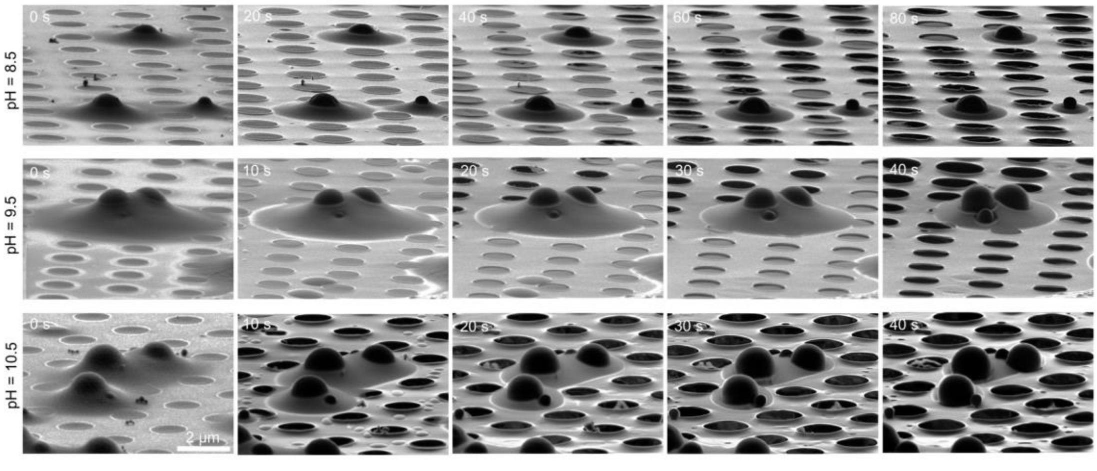


**Supplementary Fig. 13:** Time-dependent Cryo-FIB images of the droplets formed by WRWW under the gallium ion beam sputtering at different pH (WRWW, 12 mg/mL, pH 7.5)

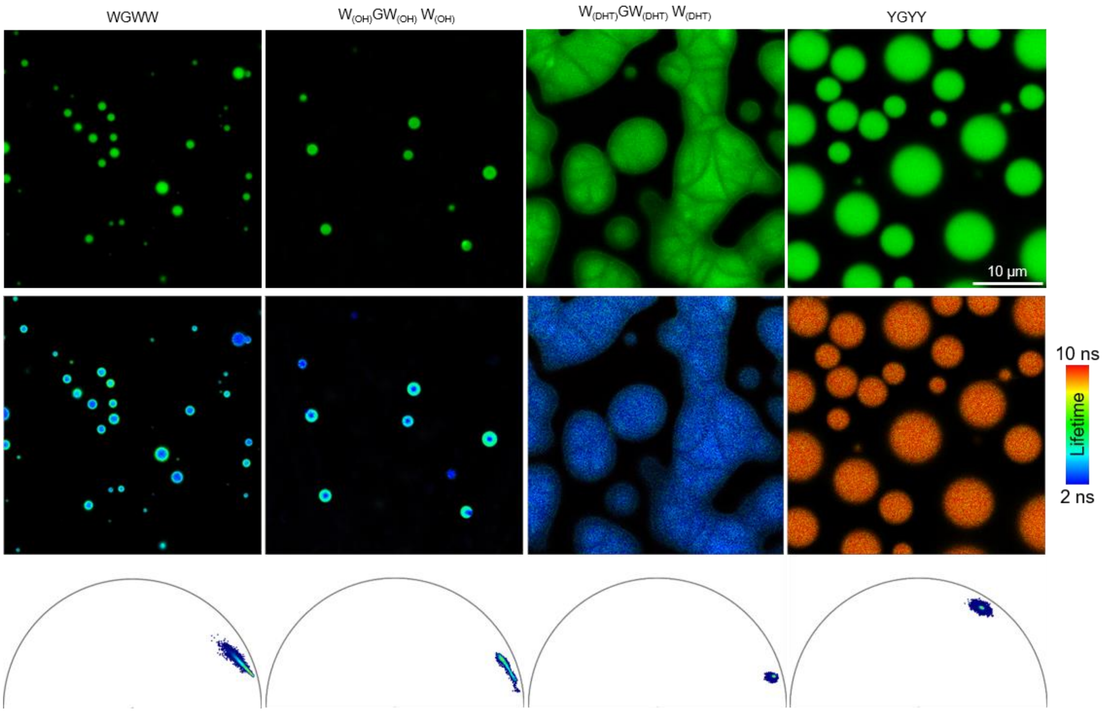


**Supplementary Fig. 14:** The confocal and fast FLIM images of the peptides WGWW (4 mg/mL, pH 7.5), $W_{(OH)}GW_{(OH)}W_{(OH)}$ (8 mg/mL, pH 7.5), $W_{(DHT)}GW_{(DHT)}$ $W_{(DHT)}$ (15 mg/mL, pH 7.5), YGYY (30 mg/mL, pH 7.5).

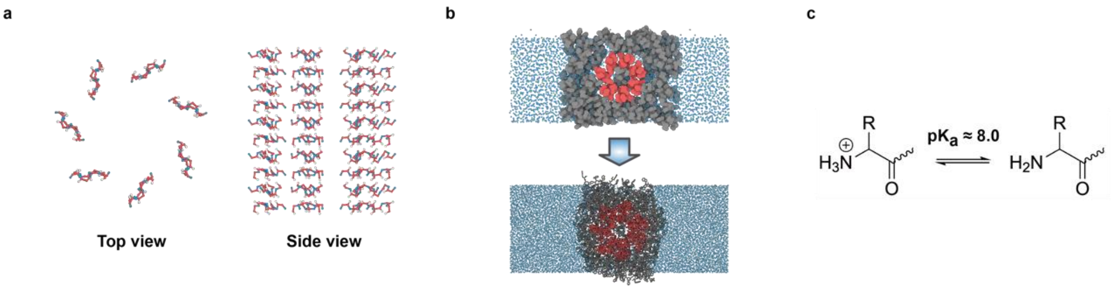


**Supplementary Fig. 15: a,** Anti-parallel structure of the fiber-like core, depicting only the backbone atoms. **b,** Multiscale simulation. Initial structure of all-atom simulation (bottom) was prepared using MARTINI simulation (top). The color coding is consistent across two simulation models: core peptides (red), shell peptides (black), and water molecules (blue). **C,** Deprotonation occurs on N-termini of peptides below pH ≈ 8.0.

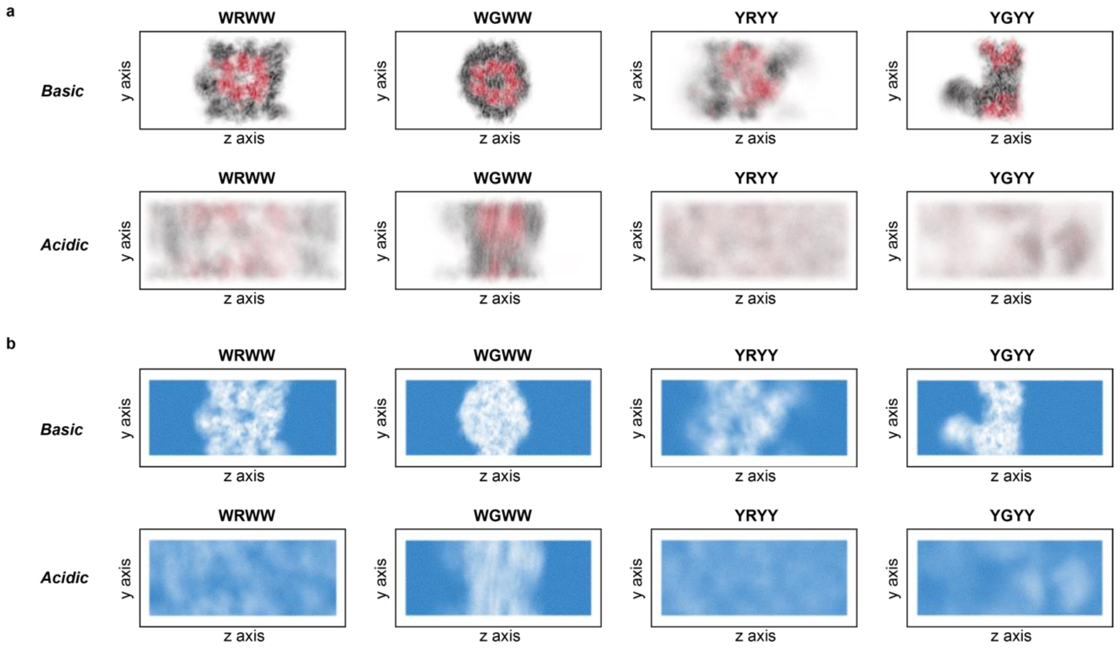


**Supplementary Fig. 16: a,** Peptide and **b,** Water atom density profile along *yz* plane of the simulation box for the last 250 ns of the simulation. The color coding is consistent across all panels: core peptides (red), shell peptides (black), and water molecules (blue). The first and second rows of each figure represent density profiles under basic and acidic conditions, respectively.

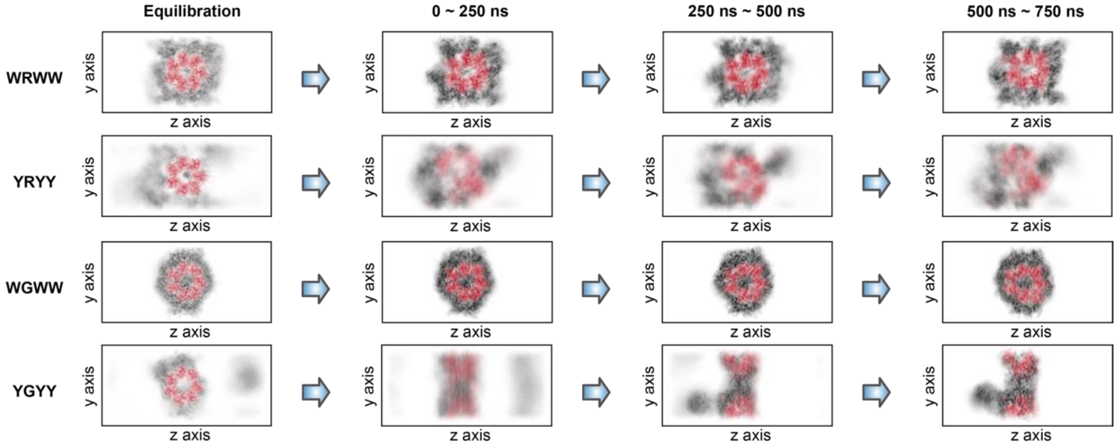


**Supplementary Fig. 17:** Average atom density profile of WRWW, YRYY, WGWW, YGYY in different simulation time windows (Basic condition). For clarity, only peptides are depicted. The color coding is consistent across all panels: core peptides (red), shell peptides (black).

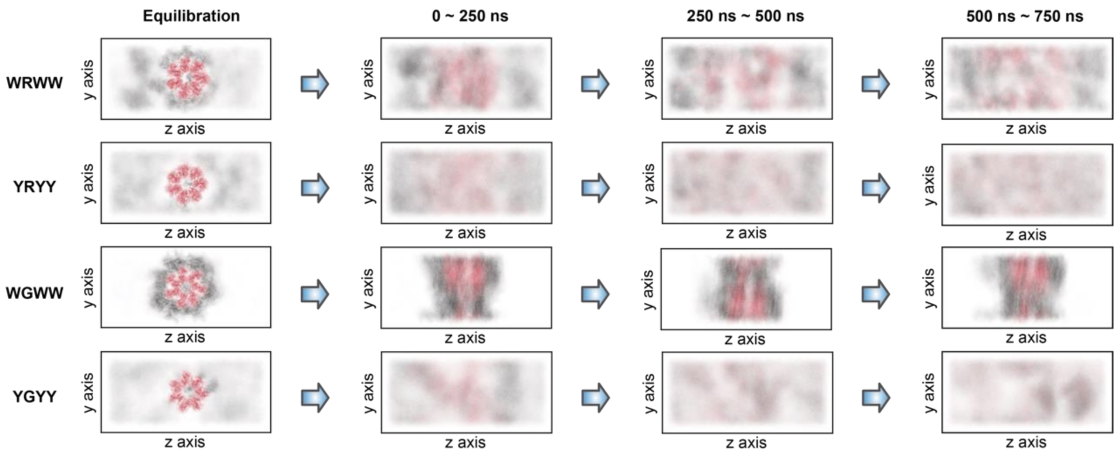


**Supplementary Fig. 18:** Average atom density profile of WRWW, YRYY, WGWW, YGYY in different simulation time windows (Acidic condition). For clarity, only peptides are depicted. The color coding is consistent across all panels: core peptides (red), shell peptides (black).

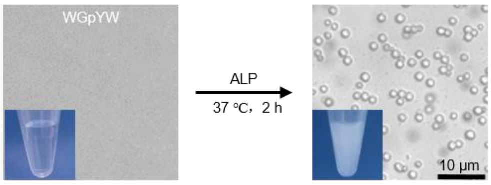


**Supplementary Fig. 19:** Enzyme controlled LLPS. WGpYW (8 mg/mL, pH 7.5), Alkaline Phosphatase（ALP) (5U/mL).

### Details of Multiscale Simulations for Tetrapeptide Droplets

#### 1. Construct anti-parallel structure as the core of condensate droplets

We began with the CyroEM structure of the 1-KMe$_3$ fiber[1], using it as a reference for the fiber-like core structure of tetrapeptide condensates. From this reference structure, we isolated a single layer, which contains ten peptide chains. We retained only the backbone atoms and removed the rest atoms. Then, using software Scwrl4[2], we added the side chain atoms based on the tetrapeptide sequences we studied experimentally: WRWW, WGWW, YRYY, and YGYY. This provided us with a single-layer structure.

Subsequently, we utilized MDTraj[3] to rotate each chain within this layer by 180°. This rotation was performed along an axis that passed through the geometric center of each chain and was also orthogonal to the plane formed by the geometric center, N-terminus, and C-terminus. The rotaion on each chain resulted in a layer oriented in the opposite direction.

Then we stacked the two layers and then duplicating the resulting structure five times, and obtained a ten-layer anti-parallel structure containing 70 peptide chains. The inter-layer distance was kept the same with the layer distance in the reference 1-KMe$_3$ structure.

The C-termini of all the tetra-peptides are capped using amides, while the N-termini remain uncapped.

### 2. Build initial configurations for multiscale simulations

Based on the fiber-core and liquid-shell model, we built initial configurations for multiscale simulations using the constructed anti-parallel tetrapeptide fiber cores. We simulated both acidic and basic environments for tetrapeptide systems including WRWW, WGWW, YRYY, and YGYY, resulting in a total of 8 simulations.
Considering the pKa of the uncapped N-terminus is approximately 8.0[4], the N-terminus of the tetra-peptides is protonated under acidic conditions. As our multiscale simulation approach started from MARTINI simulations[5], for which we used martinize2[6] to convert the all-atom structure of our constructed anti-parallel fiber cores, including single molecule peptides, into the MARTINI representations. Using martinize2, we assigned the N-terminus of each peptide with a neutral charge for simulations in basic environment, or one unit positive charge for simulations in basic environment.
The anti-parallel core was positioned in the center of a 4.9 nm × 8.0 nm × 10 nm box, with the anti-parallel core aligned along the box's *x*-axis. The *x* dimension of the box is roughly 10 times the inter-layer distance of the core structure. We then randomly inserted 130 peptides into this box, resulting in a total of 200 peptides. Subsequently, the box was expanded to dimensions of 4.9 nm × 8.0 nm × 20 nm. Finally, we solvated the box with MARTINI water and added 100 mM NaCl and neutralized the system.
In addition to above eight simulation systems, we also constructed a core-only system for WRWW in a basic environment, mirroring the setup of the previous systems but without inserting free peptides.

### 3. MARTINI simulation

We initiated the simulations with steepest descent energy minimization. Following the energy minimization, we conducted an NPT equilibration for 20 ns, using a timestep of 20 fs. The temperature of the protein and solvent was maintained at 303.15 K by individually coupling them to velocity-rescaling thermostats with a time coupling constant of 1 ps. To control pressure, we employed a Berendsen barostat with a reference pressure of 1 bar, a compressibility of $4.5\times10^{-5}$ $bar^{-1}$, and a time coupling constant of 5 ps. For the production simulations, we used an NPT ensemble with separate temperature coupling for the protein and solvent, both set to 303.15 K using velocity-rescaling thermostats with a time coupling constant of 1 ps. To couple pressure, we utilized a Parrinello-Rahman barostat with a time coupling constant of 10 ps, a compressibility of $4.5\times10^{-5}$ $bar^{-1}$, and a reference pressure of 1 bar. The production simulations lasted for 1000 ns with a timestep of 20 fs. Positional restraints of 500 $kJ\cdot mol^{-1}\cdot nm^{-2}$ were set on backbone beads of the core in the whole MARINI simulation. All the simulations were performed using GROMACS 2022[7]. For energy minimization, we utilized the double-precision version of GROMACS to solve atom clashes. For all other aspects of the simulations, we employed the mixed-precision version of GROMACS.

### 4. Atomistic simulation

We used the CHARMM36m force field[8] to describe proteins in atomistic level. We used the backward script[9] to reconstruct and optimize atomistic configurations of proteins from the last frame of MARTINI simulation trajectories. The backward script is capable of converting MARTINI configurations to atomistic representations and ensures configuration stability through a series of short energy minimization and restrained molecular dynamics simulations.

Position restraints were set on backbone and side chain atoms of the anti-parallel core structure, with a force constant of 1000 $kJ \cdot mol^{-1} \cdot nm^{-2}$ for backbone atoms, and 100 $kJ \cdot mol^{-1} \cdot nm^{-2}$ for side chain atoms, respectively.

We initiated the atomistic simulations with steepest descent energy minimization, then equilibrated the system in the NVT ensemble for 100 ps with a timestep of 1 fs. To maintain the temperature at 303.15 K, we used a Nose-Hoover thermostat, separately coupling the protein and solvent with a time coupling constant of 1.0 ps.

Next, we performed four NPT equilibrations. Each NPT equilibration lasted for 5 ns with a timestep of 2 fs. The temperature was controlled in the same manner as during NVT equilibration, and for pressure coupling, we employed a Parrinello-Rahman barostat with a compressibility of $4.5 \times 10^{-5}$ $bar^{-1}$, a reference pressure of 1 bar, and a time coupling constant of 2 ps. Throughout these four NPT equilibrations, position restraints on the core structure were gradually reduced to zero. Specifically, in four NPT equilibrations, the force constants for the backbone were decremented as 400 $kJ \cdot mol^{-1} \cdot nm^{-2}$, 200 $kJ \cdot mol^{-1} \cdot nm^{-2}$, 100 $kJ \cdot mol^{-1} \cdot nm^{-2}$, and 0. For the side chains, the force constants were decremented as 40 $kJ \cdot mol^{-1} \cdot nm^{-2}$, 20 $kJ \cdot mol^{-1} \cdot nm^{-2}$, 10 $kJ \cdot mol^{-1} \cdot nm^{-2}$, and 0.

We then performed the production simulation in the NPT ensemble for 750 ns with a timestep of 2 fs. Temperature was controlled using the Nose-Hoover thermostat, with separate coupling for the protein and solvent and a time coupling constant of 1.0 ps. Pressure control during the production simulation using a Parrinello-Rahman barostat with a compressibility of $4.5 \times 10^{-5}$ $bar^{-1}$, a reference pressure of 1 bar, and a coupling constant of 2 ps.

All the simulations were performed using GROMACS 2022. For energy minimization, we utilized the double-precision version of GROMACS to solve atom clashes. For all other aspects of the simulations, we employed the mixed-precision version of GROMACS.

### 5. Simulation analysis

The all-atom simulations were analyzed to determine the average atom density profiles. As stated above, the simulations consisted of a 140 ns equilibration simulation followed by a 750 ns production simulation. Snapshots of the simulation trajectories were saved at 1 ns intervals. The production simulation trajectory was divided into three equal time windows of 250 ns each. We analyzed the average atom density for both the equilibration simulation and the three segments of the production simulation, separately. The average atom density was calculated across the *y*-z plane of the simulation box, which is orthogonal to the fiber-like core. This *y-z* plane was divided into a grid with a resolution of 0.1 nm by 0.1 nm. The average number of atoms within each grid cell across the simulation time windows was computed. Peptides in the core, peptides in the shell, and water molecules were quantified separately. The analysis was performed using the GROMACS and MDTraj software packages.
